# Nuclear GAPDH in cortical microglia mediates stress-induced cognitive inflexibility

**DOI:** 10.1101/2022.06.21.497065

**Authors:** Adriana Ramos, Koko Ishizuka, Ho Namkung, Lindsay N. Hayes, Atsushi Saito, Arisa Hayashida, Rupali Srivastava, Noah Elkins, Trexy Palen, Elisa Carloni, Tsuyoshi Tsujimura, Coleman Calva, Satoshi Ikemoto, Rana Rais, Barbara S. Slusher, Minae Niwa, Toshiaki Saitoh, Eiki Takimoto, Akira Sawa

## Abstract

We report a mechanism that underlies stress-induced cognitive inflexibility at the molecular level. In a mouse model under subacute stress in which deficits in rule shifting tasks were elicited, the nuclear glyceraldehyde dehydrogenase (N-GAPDH) cascade was activated specifically in microglia in the prelimbic cortex. The cognitive deficits were normalized with a pharmacological intervention with a compound (the RR compound) that selectively blocked the initiation of N-GAPDH cascade without affecting glycolytic activity. The normalization was also observed with a microglia-specific genetic intervention targeting the N-GAPDH cascade. Furthermore, hyperactivation of the prelimbic layer 5 excitatory neurons, which are known to be a neuronal substrate of cognitive inflexibility, was also normalized by the pharmacological and microglia-specific genetic interventions. The RR compound may offer a mechanism-driven, translational opportunity against stress-induced cognitive inflexibility. Taken together, we show a pivotal role of cortical microglia and microglia-neuron interaction in stress-induced cognitive inflexibility. We underscore the N-GAPDH cascade in microglia, which causally mediates stress-induced cognitive alteration.

## INTRODUCTION

Cognitive flexibility is a construct of executive function that adapts our decision-making to new experiences [1]. This behavioral construct is closely related to creativity and empathy, enabling us to not only adapt, but also to think outside of the box, taking other perspectives into consideration [1]. Cognitive flexibility is predominantly regulated by the prefrontal cortex [2–4]. This function is sensitive to, and negatively affected by conditions of stress [5–7]. Accordingly, cognitive inflexibility is frequently observed in an array of brain disorders, including schizophrenia (SZ) and Alzheimer’s disease (AD) [8–10]. In these conditions, impairment of cognitive flexibility directly leads to functional deficits and worsens clinical outcomes for patients [11,12]. Nevertheless, pathophysiological mechanisms for this impairment at the molecular and cellular levels remain to be elucidated.

At the molecular level, inflammation and redox imbalance have been frequently associated with poor cognitive performance [13–22]. Biofluids from patients with cognitive deficits have shown molecular signatures of inflammatory processes and excess oxidative stress [17,20,23–25]; however, it is unknown how such molecular changes are related to specific symptoms at the mechanistic level. Meanwhile, basic neurobiology has shown that neuroinflammatory molecules can modulate synaptic structures and functions in rodent models [18,19]. An array of rodent models with cognitive deficits have been used to connect the gaps between basic neurobiology and clinical observations [26–29]. Some models are generated by administering lipopolysaccharide (LPS), polyinosinic:polycytidylic acid (poly I:C), ethanol, or cuprizone [30–34]. Repeated administration of low-dose LPS consistently shows excess oxidative stress and prolonged changes in microglia [35–37]. Cognitive performance has also been precisely addressed in this model after resolution of the transient sickness behavior that occurs upon LPS injection [38].

Among representative stress-triggered intracellular signaling, we have studied a cascade in which glyceraldehyde-3-phosphate dehydrogenase (GAPDH) acts as a sensor of cellular stress [39,40]. Beyond its classic role in glycolysis [41,42], during stress, a pool of this multi-functional protein [40,43] is post-translationally modified at specific residues, such as cysteine 150 (Cys-150) and lysine 160 (Lys-160), which converts this glycolytic enzyme into a signaling molecule [44]. A wide range of stressors that ultimately lead to oxidative/nitrosative stress in the cells can trigger this cascade, which include dexamethasone for thymocytes and LPS for macrophages [44,45]. At the molecular levels such stressors lead to *S*-nitrosylation (or possibly oxidation) at Cys-150, which allows post-translationally modified GAPDH to interact with a chaperon protein Siah1 creating a protein complex of GAPDH-Siah1. Through this process, GAPDH loses its glycolytic activity and is translocated to the nucleus as a GAPDH-Siah1 complex [44]. The nuclear translocation of a pool of GAPDH implies a gain-of function that has a significant impact on cellular signaling, whereas a loss-of-function of 1-2% of the total glycolytic activity due to this conversion is negligible on overall cell metabolism [44]. Accordingly, GAPDH-Siah1 protein binding is a key regulatory process that initiates the nuclear-GAPDH (N-GAPDH) cascade, which can be selectively blocked by several compounds without affecting the glycolytic activity of GAPDH. These selective blockers include structural analogues of deprenyl (e.g., CGP 3466B) [46]. We have recently reported that (1R, 3R)-1, 3-dimethyl-2-propargyl-1, 2, 3, 4-tetrahydroisoquinoline (designated as RR in this study) is a deprenyl structural analogue with the highest potency among the analogues, as far as we are aware, for blocking the GAPDH-Siah1 protein interaction [47]. Our group also reported that the replacement of lysine 225 for alanine (K225A) on GAPDH abolished GAPDH-Siah1 binding [44]. Although this cascade has been mainly studied in the context of cell death [48], activation of the N-GAPDH cascade was also reported in animal models with behavioral deficits that do not accompany cell death [49]. Nevertheless, the mechanism(s) whereby the N-GAPDH cascade mediates stress-induced behavioral alteration remain elusive.

Here we used a mouse model with repeated administration of low-dose LPS [<u>l</u>ow-dose <u>s</u>tressors (LPS) at <u>m</u>ultiple times: LSM model] in which we observed stress-induced cognitive inflexibility together with selective activation of the N-GAPDH cascade in cortical microglia. The cognitive deficits were reverted with pharmacological intervention or microglia-specific genetic intervention targeting the initial steps of the N-GAPDH cascade. Hyperactivation of the prelimbic excitatory neurons, known as a key neural substrate of the deficit in rule shifting tasks, was also normalized with the same set of pharmacological or microglia-specific genetic intervention to the N-GAPDH cascade. Starting from a specific question of the N-GAPDH stress cascade, we now demonstrate a molecular/cellular mechanism of stress-induced cognitive inflexibility, a general question in neurobiology and biological psychiatry. A stress-induced signaling cascade (the N-GAPDH cascade) specifically activated in microglia mediates the cognitive inflexibility via a prefrontal microglia-neuron crosstalk.

## METHODS

### Mice

Male C57BL/6J, CMV-Cre, Cx3cr1-CreER, VGlut1-Cre, VGAT-Cre, Thy1-GCaMP6s, GCamMP6 floxed, and K225A-GAPDH floxed [47] mice were used for the experiments. All mice were on a C57BL/6 background.

K225A-GAPDH floxed model. Genetically modified mice in which lysine 225 (K) responsible for the binding between GAPDH and Siah1 was replaced with an alanine (A) residue, were generated using a homologous recombination approach [47]. In this mouse model GAPDH is not able to translocate to the nucleus, therefore the GAPDH nuclear pathway is blocked [50]. These animals were crossed with Cx3cr1-CreER promoter line in order to target microglia.

Eight to ten weeks old mice were used for all experiments. Mice were housed (5 mice maximum per cage) and maintained under a 12-h light: 12-h dark cycle, at a constant temperature and humidity (20-24°C, 35-55%), with food and water available *ad libitum*. The Institutional Animal Care and Use Committee at Johns Hopkins University approved all protocols involving mice that were used in this study.

### Chemicals

LPS and RR were reconstituted with saline buffer. Mice were injected with saline or RR (0.25 mg/kg/day i.p for 5 days) 1 day prior to a low-dose LPS treatment at multiple times (1.5×10^5^ EU/kg/day i.p for 4 days) (LSM) and were co-treated with RR (or saline) and LPS for the following 4 days. Tamoxifen was used to drive the Cx3cr1-CreER promoter. The tamoxifen solution (10 mg/mL) was freshly prepared by sonicating tamoxifen freebase in sunflower seed oil (S5007, Sigma-Aldrich) at room temperature for 10-12 min (with intermittent 20 sec vortexing every 3-4 min). This solution was stored at −20°C for several months. Mice aged 4-5 weeks were administered tamoxifen by oral gavage at a dosage of 100 mg/kg body weight, once a day for two consecutive days. As microglia were the target cell, all experiments were performed 5 weeks after the last tamoxifen injection.

### RR pharmacokinetic analysis

Vehicle control or RR treated mice (0.25mg/kg/day i.p. for 5 days) were euthanized 1h after the last injection (n=5 mice). Blood samples were collected in heparinized tubes by cardiac puncture and centrifuged (3000 × g for 10 min) to obtain plasma. Brains were dissected and both plasma and brains were stored at −80°C until bioanalysis. RR compound levels in the plasma and brain were measured using high-performance liquid chromatography with tandem mass spectrometry (LC/MS-MS). Briefly, standards were prepared by spiking the compound in naïve mouse plasma or tissue from 0.003-100 nmol/mL or nmol/g. Standards, plasma (20 μL) or weighed brain samples were placed in low retention microcentrifuge tubes and extracted using methanol containing losartan as an internal standard (5 μL methanol/mg tissue). Samples were vortex mixed and centrifuged (16,000 × g for 5 min at 4°C). The supernatants (80 μL) were transferred to a 96 well plate and 2 μL was injected for analysis. Samples were analyzed on an UltiMate 3000 UHPLC coupled to a Q Exactive Focus orbitrap mass spectrometer (Thermo Fisher Scientific Inc., Waltham MA). Samples were separated on an Agilent EclipsePlus C18 RRHD (1.8 μm) 2.1 × 100 mm column. The mobile phase consisted of water + 0.1% formic acid (A), and acetonitrile + 0.1% formic acid (B). Separation was achieved at a flow rate of 0.4 mL/min using a gradient run. Quantification was performed in product-reaction monitoring (PRM) mode (RR 200.1434> 145.1, 82.0650) using Xcalibur software.

### Behavioral assays

The Institutional Animal Care and Use Committee at Johns Hopkins University approved all protocols used in this study.

#### Cognitive flexibility tasks

We followed a protocol described by Cho *et al* [3] with some adaptations from Heisler *et al* [51]. Briefly, mice were single-housed and habituated to a reverse light/dark cycle at least two weeks before starting the task. Food intake was restricted 5 days before starting, corresponding to the period in which the mice were adapted to the bowls and reward. At the start of each trial, the mouse was placed in its home cage to explore two bowls, each containing one odor and one digging medium, until it dug in one bowl, signifying a choice. The bait was a piece of a peanut butter chip (approximately 5-10 mg in weight) and the cues, either olfactory (odor) or somatosensory and visual (texture of the digging medium which hides the bait), were altered and counterbalanced. All cues were presented in small animal food bowls that were identical in color and size. Digging media was mixed with the odor (0.1% by volume) and peanut butter chip powder (0.1% by volume). All odors were ground dried spices (McCormick garlic and coriander). For each trial, while the odor-medium combination present in each of the two bowls may have changed, the stimulus (e.g., a particular odor or medium) that signaled the presence of the food reward remained constant over each portion of the task (initial association, rule shift). If the initial association paired a specific odor with the food reward, then the digging medium would be considered the irrelevant dimension. The mouse was considered to have learned the initial association between stimulus and reward if it made 8 correct choices during 10 consecutive trials. Following the initial association, the rule-shifting portion of the task began, and the stimulus associated with the reward underwent an extra-dimensional shift. The mouse was considered to have learned the extra dimensional rule shift if it made 8 correct choices during 10 consecutive trials. In all portions of the task, after the mouse made an error on a trial or 3 min passed with no choice, the mouse was transferred to the holding (punishment) cage for a minimum of 60 sec. If the animal made 6 consecutive no choices (3 min without a choice) this was recorded as a failure to participate.

#### Forced swim test

Each mouse was put in Plexiglass cylinder tanks measuring 19 cm in height and 13 cm in diameter with a water level of 16 cm. Mice were scored for immobility time over a 6-min uninterrupted test session.

#### Open field test

An individual mouse was placed near the wall-side of a 50 × 50 cm open-field arena. The movement of the mouse was recorded by laser beam breaks for 30 min using an infrared activity monitor (San Diego Instruments). The data was automatically analyzed with PAS-Open Field software (San Diego Instruments). Time in the center of the field and total activity were measured. The open field arena was cleaned with 70% ethanol and wiped with paper towels between each trial.

### Cell biology, biochemistry, and histology

#### Microglia and astrocytes sorting

CD11b+ or ACSA2+ beads were used to isolate microglia or astrocytes, respectively, by magnetic sorting. The “rest” fraction corresponded with the negative fraction isolated after separating microglia and astrocytes. Microglia were isolated by fluorescence-activated cell-sorting (FACS) using CD45, CD11b, and p2ry12 antibodies (Biolegend). These cells were directly sorted into extraction lysis buffer for qPCR. RNA extraction was performed using the Arcturus Pico Pure RNA isolation kit (Thermo Fisher).

#### Immunoblotting

Microglia, astrocytes, or rest cells were lysed in CHAPS buffer (50 mM Tris, 150mM NaCl, 1% SDS, and 1% Triton X-100) containing a proteinase inhibitor and phosphatase inhibitor cocktails. Six rounds of sonication were performed as follows: 15 sec on, 15 sec off, and 40% amplitude with a Q125 Sonicator (Q Sonica Sonicators). Lysates were centrifuged for 2 min at 10,000 rpm, the supernatant was collected, quantified using the Pierce BCA Protein assay (Thermo Scientific), aliquoted, and stored at −80°C until use. Lysates were thawed and 4× NuPAGE LDS sample buffer (Thermo Scientific) and 10% β-mercaptoethanol (Sigma Aldrich) were added. Lysates were heated to 95°C for 10 min and run on NuPAGE 4%-12% bis-tris gels (Invitrogen). Proteins were transferred to PVDF membranes (BioRad) and blocked for 1 h at RT in 5% BSA in Tris buffered saline with 0.05% Tween 20 (TBST). Membranes were incubated in primary antibody overnight at 4°C, washed and incubated in secondary antibody HRP for 1 h at RT. After washing, membranes were developed using either the Super Signal West Dura or Super Signal West Femto Chemiluminescent Substrate (Thermo Scientific).

#### Immunoprecipitation (IP)

200 μg of total protein was used for Siah1 IP, the total volume was increased to 1ml with supplemented CHAPS buffer, 1 μg of antibody was added and incubated O/N at 4°C with continuous agitation. On the next day cell lysates were incubated for 2 h at 4°C with 30 μL of TrueBlot Anti-Goat or Rabbit Ig beads (Rockland laboratories).

#### Dot blot

The potency of RR and other deprenyl derivatives to block the binding between GAPDH and Siah1 was evaluated using dot blot. A mixture of GAPDH and Siah1 recombinant proteins was incubated O/N at 4°C, with each deprenyl derivative (2 nM) or deprenyl (2 nM) or empty solution. The possible protein complexes were immunoprecipitated using a Siah1 antibody as described above, and the immunoprecipitates were loaded onto a nitrocellulose membrane. The membrane was washed and immunoblotted with a GAPDH antibody as described above.

#### Immunohistochemistry

After perfusion, mice were fixed with 4% paraformaldehyde (PFA) and brains were kept in PFA O/N at 4°C. The following day brains were transferred to a solution of phosphate buffered saline (PBS) containing 30% sucrose for 1-2 days, then 12-30-μm cryosections were collected on glass slides. Sections were blocked in TBS containing 0.5% Triton (TBS-T) and 10% normal donkey serum (NDS) for 2 h at RT. Sections were then incubated with primary antibodies and kept at 4°C for 24-48 h. Subsequently, sections were washed three times with TBS-T for 5 min each and incubated with fluorescently conjugated-secondary antibodies for 2 h at RT. After three washes with TBS-T, sections were mounted and coverslipped. Fluorescence images were obtained using a Zeiss LSM 700, Olympus FV1000.

### Physiological experiments

#### Acute brain slice preparation

Mice were anesthetized with ether, perfused using cold N-methyl-D-glucamine (NMDG) based cutting solution containing (in mM): 135 NMDG, 1 KCl, 1.2 KH_2_PO_4_, 1.5 MgCl_2_, 0.5 CaCl_2_, 10 Dextrose, 20 Choline Bicarbonate, (pH 7.4) as previously described [52]. Brains were dissected and mounted on a vibratome (Leica VT100S). Cortical slices 250 μm thick were cut in ice-cold N-methyl-D-glucamine (NMDG) based cutting solution. Cortical slices were then transferred to artificial cerebral spinal fluid (ACSF) containing (in mM): 119 NaCl, 2.5 KCl, 2.5 CaCl_2_, 1.3 MgCl_2_, 1 NaH_2_PO_4_, 26.2 NaHCO_3_, and 11 Dextrose (292–298 mOsm/L) and were maintained at 37° C for 40 min, and at room temperature thereafter. Both the NMDG solution and ACSF were bubbled continuously with 95% O_2_/5% CO_2_. All slice imaging experiments were carried out at room temperature.

#### Slice Ca imaging

Fluorescence dynamics arising from GCaMP6s VGlut1, VGAT, or Thy1 positive neurons were recorded using a fixed-stage microscope (BX61-WI, Olympus) with optical lens (40×-0.80, LumPLanFL N), 300 W Xenon lamp using a GFP filter, high speed galvo mirror with filters (DG4-1015, Shutter Instrument Company), and an infrared-sensitive CCD camera (iXon3, ANDOR Technology). Images were processed with digital imaging software (MetaFluor® for Olympus and Metamorph Advanced Molecular Device). All these systems were on an anti-vibration floating table (Technical Management Corp.) and connected to a PC (Windows 7, Microsoft). For imaging analysis, raw videos were pre-processed by applying ×4 spatial down-sampling to reduce file size and processing time, but no temporal down-sampling was applied [53]. To reliably deal with the large fluctuating background, we applied the CNMFe algorithm that is a recent adaptation of the CNMF algorithm [54], enabling us to identify individual neurons, obtain their fluorescent traces, and deconvolve fluorescence signals into neural activity. To define the initial spatial components, candidate seed pixels were manually selected from peak-to-noise (PNR) graphs of the field of view (FOV) [55]. Calcium events with z-scores < 8 or those that did not have a >0.5 AUC were excluded from analyses because events of this magnitude did not reliably retain transient, calcium-event characteristics across animals.

### Statistical analysis

All data are expressed as the mean ± SEM. Quantification of confocal microscopy, immunoblots, Ca^2+^ imaging, and slice electrophysiology recordings was performed using Imaris, ImageJ, and MATLAB, and Clampfit respectively. Statistical differences between two groups was determined by a two-tailed unpaired Student’s *t-*test. For multiple group datasets, two-way or one-way ANOVA analysis was used, followed by Tukey’s multiple comparisons test for more than two groups. For multiple group datasets of slice electrophysiology recordings, repeated measures ANOVA analysis was used, followed by Dunnett’s post-hoc tests. Statistical tests used to measure significance are indicated in each figure legend along with the corresponding significance level (p value).

## RESULTS

### The N-GAPDH cascade underlies cognitive inflexibility in a stress-induced mouse model

To test whether the N-GAPDH cascade plays a mechanistic role linking stress and behavior at the molecular level, we used the LSM model (**Fig. 1a**). We observed an excess of oxidative stress and activation of the N-GAPDH cascade in the brains of LSM mice (**Supplementary Fig. 1a, b**). In contrast, we did not observe any sign of blood brain barrier damage, nor cell death in the brain (**Supplementary Fig. 1c-f**). Signs of sickness behavior were not present 24 h after treatment [38], when all biochemical or behavioral outcomes were recorded. No changes in locomotor and swimming activity were observed in LSM mice compared to control mice (**Supplementary Fig. 1g, h**). However, LSM mice presented significant deficits in cognitive flexibility as measured by the attentional set-shifting task, but showed no impairment in learning the initial association portion of the task (**Fig. 1a-c and Supplementary Fig. 1i**). Moreover, we observed that LSM mice performed more perseverant errors than control mice (**Fig. 1d**). These results allowed us to conclude that LSM mice were less flexible, requiring more attempts in order to disengage from the paradigm that they learned during the initial association.

**Fig. 1.**
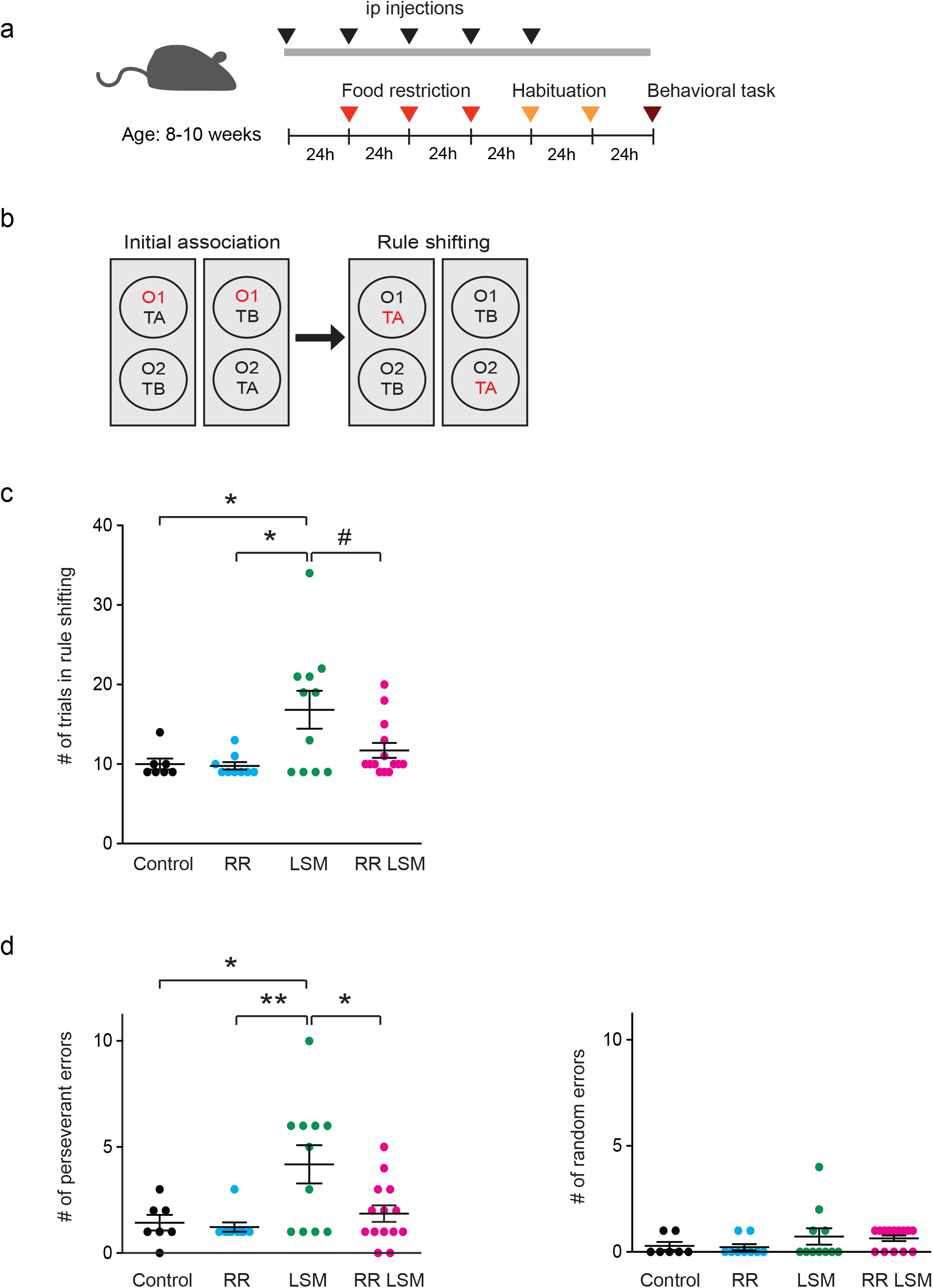
The N-GAPDH cascade underlies stress-induced cognitive inflexiblity in LSM mouse model. **a**) Schematic diagram of the mouse paradigm and behavioral task. Eight to ten week-old mice were adapted to a reverse light/dark cycle. Drug injections were made intraperitoneally (ip). Mice were food restricted for 72 h before starting habituation, then habituated for 2 consecutive days to the bowls, digging media, and food reward before testing began. Mice that reached criteria were tested with the behavioral task on the following day. Black arrowhead, injection; red line, calorie restriction; yellow arrowhead, habituation; and brown arrowhead, behavioral test. (**b**) Schematic diagram illustrating the rule-shifting task. Two bowls were presented to mice, each baited by an odor (O1 or O2) and a textured digging medium (TA or TB). Mice had to find a food reward that was associated with a particular stimulus (indicated in red) buried in the digging medium. Mice first learned an initial association; once mice reached the learning criterion (eight correct out of ten consecutive trials), this association underwent an extradimensional rule shift. (**c**) Performance of Control, RR, LSM, and RR LSM mice in the rule shifting task. Each dot represents data from an individual mouse. Two-way ANOVA (RR × LSM) revealed a significant main effect of RR (F_(1,37)_ = 4.68, p=0.037), a significant effect of LSM (F_(1,37)_ = 7.33, p=0.010), and a non-significant interaction between factors (F_(1,37)_ = 2.55, p=0.119). Error bars represent the mean ± SEM. ^#^p =0.051 *p <0.05 by two-way ANOVA with Tukey’s multiple comparisons test. (**d**) LSM mice performed significantly more perseverant than random errors during the ruleshifting task. Each dot represents data from an individual mouse. Two-way ANOVA (RR x LSM) revealed a significant main effect of RR (F_1,37)_ = 6.66, p=0.014), a significant effect of LSM (F_(1,37)_ = 7, p=0.012), and a non-significant interaction between factors (F(_1,37_) = 3.12, p=0.086). Error bars represent the mean ± SEM. *p < 0.05, **p < 0.01 by two-way ANOVA with Tukey’s multiple comparisons test.

To test whether the N-GAPDH cascade is involved in the regulation of this cognitive construct, we used the RR pharmacological compound to specifically block activation of the N-GAPDH cascade by interfering with the binding of GAPDH and Siah1, without affecting the glycolytic activity of GAPDH and monoamine oxidase. We confirmed *in vitro* that this compound blocked the GAPDH-Siah1 protein interaction more potently than other drug analogs (**Supplementary Fig. 2a**). In addition, we demonstrated that RR reached the brain when it was systemically injected, and even displayed a unique enrichment in the brain (**Supplementary Fig. 2b**). Therefore, we systemically administered the RR compound in LSM mice to evaluate its impact on behavioral flexibility. We confirmed that the initiation of the N-GAPDH cascade in the brain was blocked by the systemic injection of RR (RR LSM mice) (**Supplementary Fig. 1b**). Interestingly, the RR injection also successfully blocked the behavioral deficits seen in the rule-shifting paradigm. In particular, RR significantly reduced the number of perseverant errors (**Fig. 1c, d**). These results indicate that the N-GAPDH cascade is activated in response to LSM treatment and underlies cognitive inflexibility.

### Stress-induced upregulation of the N-GAPDH cascade occurs selectively in microglia of LSM mouse model

Since activation of the N-GAPDH cascade was responsible for the behavioral inflexibility deficits observed in LSM mice, we decided to determine the cellular specificity of the N-GAPDH activation. We focused on microglia and astrocytes because they are the main cell types involved in the provocation and resolution of inflammatory and oxidative stress related processes in the brain [56–58]. Interestingly, we observed a significant upregulation of N-GAPDH only in microglia, as reflected by an increased binding of GAPDH and Siah1 in microglia (**Fig. 2a,b**). A specific antibody against sulfonation of GAPDH at cysteine-150 (SulfoGAPDH Ab) [44,50] also confirmed activation of this cascade in microglia in the medial prefrontal cortex (**Fig. 2c and Supplementary Fig. 3a**), but not in astrocytes or neurons. Nuclear fractionations proved the presence of GAPDH in the nuclei of microglia (**Supplementary Fig. 3b**). Furthermore, we observed activation of this cascade in cortical but not striatal microglia (**Fig. 2d**). As expected, systemic administration of the RR compound blocked activation of the N-GAPDH cascade in microglia (**Fig. 2e**).

**Fig. 2.**
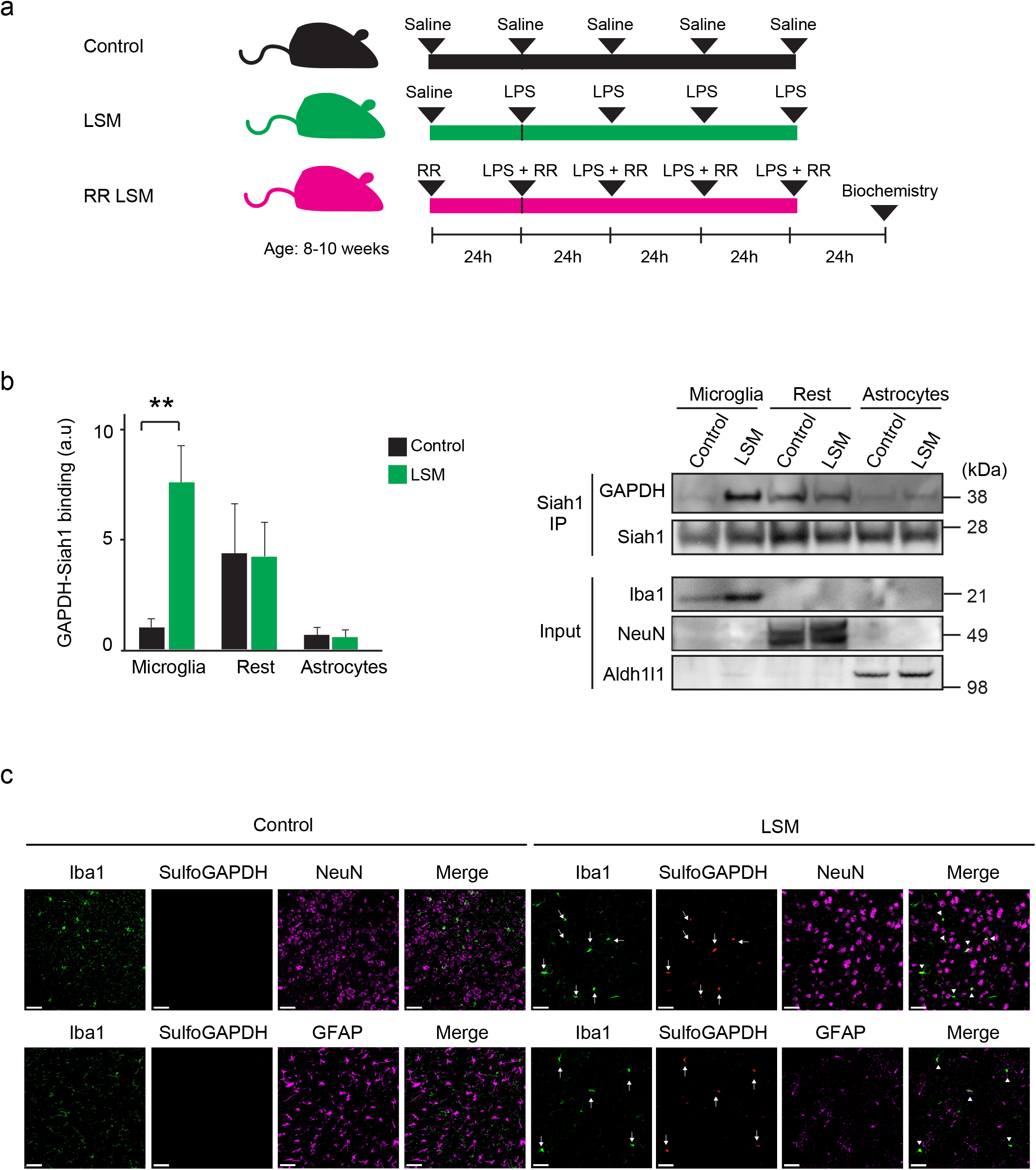
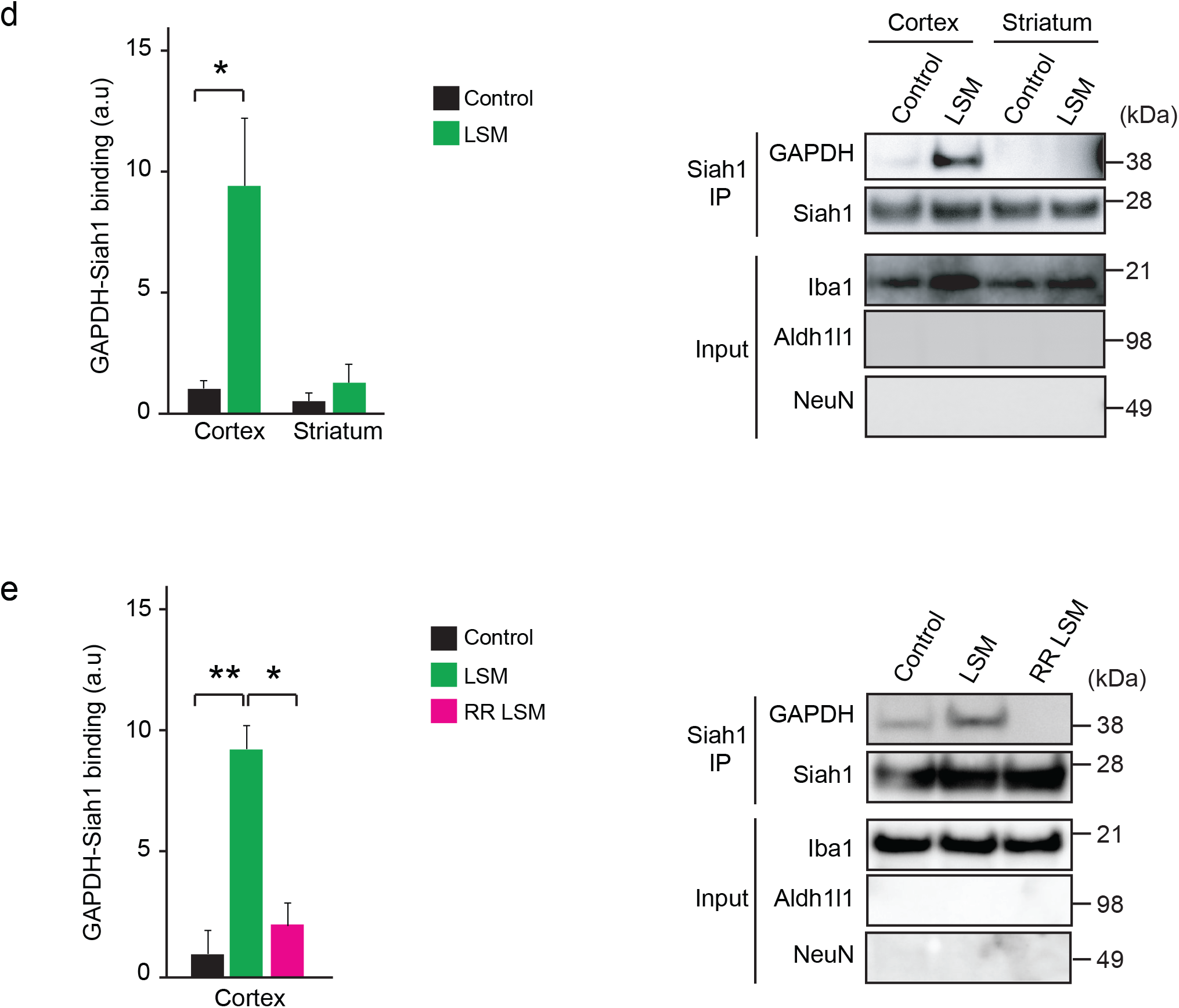
The N-GAPDH cascade is selectively upregulated in microglia of LSM mouse model. (**a**) Schematic diagram showing the mouse intraperitoneal injection paradigm used. (**b**) Levels of GAPDH-Siah1 binding (a key indicator of the N-GAPDH cascade activation). Lysates were from cortical microglia and astrocytes isolated from Control and LSM mice using magnetic sorting. “Rest” corresponds to the negative fraction after isolation of microglia and astrocytes. The purity of isolated cells was assessed by using anti-Iba1, NeuN, and Aldh1L1 antibodies that specifically recognize microglia, neurons, and astrocytes, respectively. The Y axis depicts the level of GAPDH-Siah1 binding, which was normalized by the level of Siah1. (**c**) Representative confocal images depicting sulfonated GAPDH at cysteine-150 (SulfoGAPDH) co-localized with microglia (Iba1), and not astrocytes (GFAP) or neural cells (NeuN), in LSM mice. Arrows indicate Iba1 and SulfoGAPDH signals, whereas arrowheads indicate colocalization of the Iba1 and SulfoGAPDH signals. Scale bar: 40 μm. (**d**) Level of GAPDH-Siah1 binding. Lysates were from cortical and striatal microglia isolated from Control and LSM mice. The Y axis depicts the level of GAPDH-Siah1 binding, which was normalized by the level of Siah1. (**e**) Level of GAPDH-Siah1 binding. Lysates were from cortical microglia isolated from Control, LSM, and RR LSM mice. The Y axis depicts the level of GAPDH-Siah1 binding, which was normalized by the level of Siah1. Error bars represent the mean ± SEM. *p <*p < 0.01 by unpaired two-tailed *t*-test for panel (**b**) and (**d)**, and one-way ANOVA with Tukey’s multiple comparisons test for panel (**e**). Immunoblots are representative of 5 independent experiments from cells isolated from 10 mice per group [(**b**), (**d**), and (**e**)]. Confocal images are representative of 3 mice per group (**c**).

### The N-GAPDH cascade in microglia mediates stress-induced cognitive inflexibility (deficits in a rule-shifting behavior) in LSM mouse model

In addition to the pharmacological intervention by RR, we generated a conditional mouse model designed to disrupt GAPDH-Siah1 binding and thus prevent activation of the N-GAPDH cascade under stressors [47]. We previously published that a point mutation that replaces lysine 225 of GAPDH with alanine (K225A) can abolish the GAPDH-Siah1 binding without affecting other key features of GAPDH, including its glycolytic activity [44,47]. To specifically block the N-GAPDH cascade in microglia in the central nervous system, we generated GAPDH-K225A^Cx3cr1-CreER^ knockin mice (K225A-MG mice). As expected, we did not observe an upregulation of the N-GAPDH cascade in these knockin mice even when these mice were repeatedly treated with LPS (K225A-MG LSM) (**Fig. 3a**). Further, this microglia-specific genetic manipulation to block the N-GAPDH cascade successfully prevented the behavioral deficits in the rule-shifting paradigm including a reduced number of perseverant errors in K225A-MG LSM mice (**Fig. 3b, c**).

**Fig. 3.**
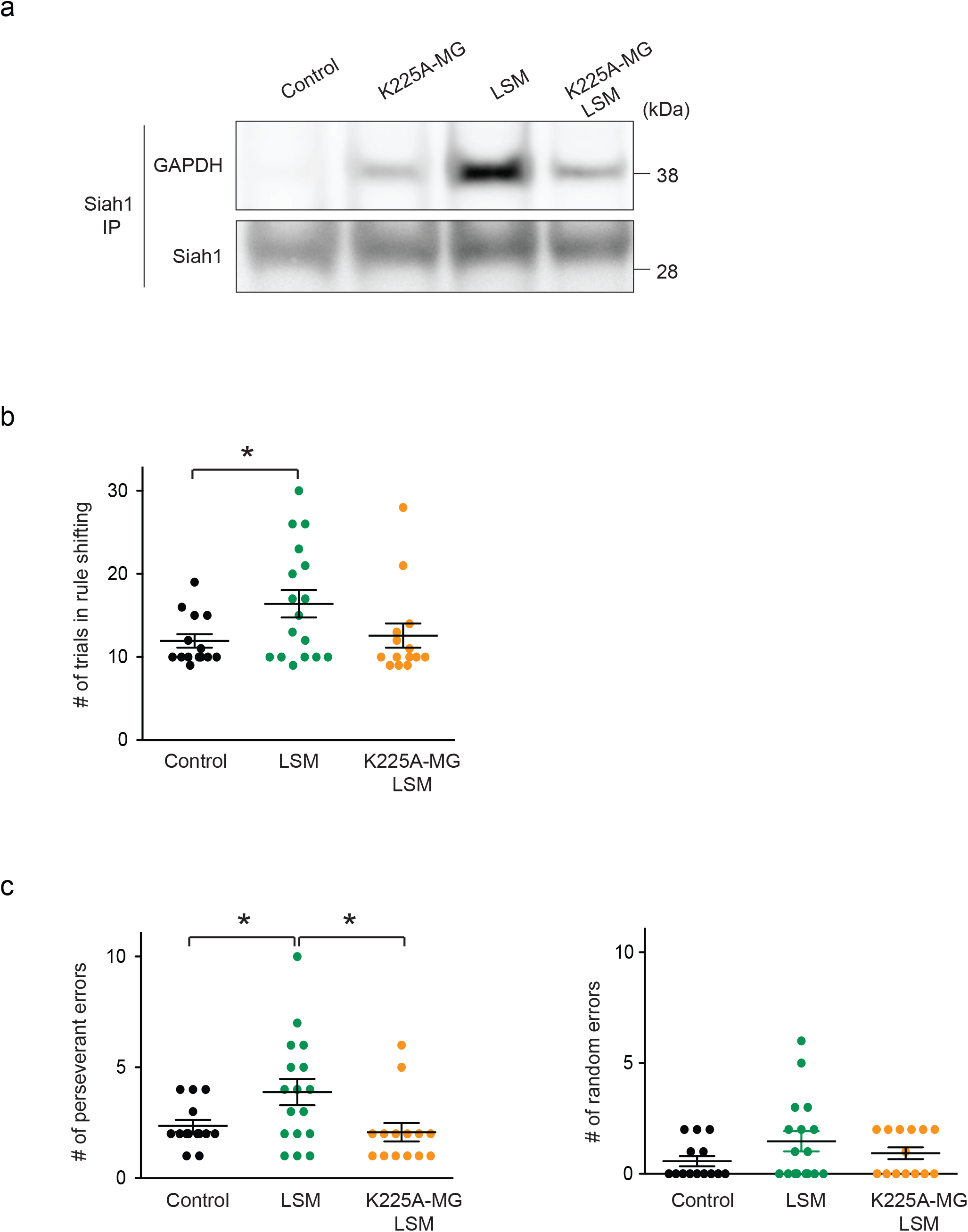
Microglia-specific genetic intervention to the N-GAPDH cascade prevents behavioral deficits in the rule-shifting paradigm in LSM mouse model. (**a**) Level of GAPDH-Siah1 binding for cortical microglia. Siah1 immunoprecipitations and immunoblots for GAPDH in Control [Cx3cr1-CreER; WT-GAPDH]; K225A-MG, [Cx3cr1-CreER; K225A-GAPDH]; LSM (LSM on control background); and K225A-MG LSM mice. (**b**) Performance of Control, LSM, and K225A-MG LSM mice in the rule shifting task. Each dot represents data from an individual mouse. (**c**) LSM mice performed significantly more perseverant than random errors during the ruleshifting task. Each dot represents data from an individual mouse. Error bars represent the mean ± SEM. *p < 0.05 by one-way ANOVA with Tukey’s multiple comparisons test for panels (**b**) and (**c**).

### The N-GAPDH cascade in microglia mediates hyperactivation of the prelimbic (PrL) layer 5 (L5) excitatory neurons in LSM mouse model

Hyperactivation of the PrL excitatory neurons is known as a key neural substrate of the deficit in rule shifting tasks [3]. We next tested whether the hyperactivation occurred in LSM mouse model, which may possibly be normalized by the pharmacological and microglia-specific genetic intervention to the N-GAPDH cascade. To address this question, we used mice that selectively expressed GCaMP6s in VGlut1-positive excitatory neurons and specifically focused on the medial prefrontal cortex (the PrL area) [3]. To assess the spontaneous activity of individual excitatory neurons, we used a constrained non-negative matrix factorization (CNMF) algorithm optimized for slice imaging [54] that allowed us to determine the frequency and amplitude of calcium events (**Fig. 4a**). Calcium transients in PrL excitatory neurons were more frequent in LSM compared with control mice (**Fig. 4b**), whereas amplitude levels were similar (**Fig. 4b**). These data indicate a plausible increase in neuronal excitability and an overall cytosolic calcium overload in LSM excitatory neurons. As shown before, no neuronal cell death was observed in LSM mice (**Supplementary Fig. 1f**). Treatment of LSM mice with RR ameliorated the increased frequency of calcium transients, suggesting a causal involvement of microglial activation of the N-GAPDH cascade (**Fig. 4b**). The causal role of microglial N-GAPDH cascade was directly shown by no increase in calcium transients frequency in K225A-MG LSM mice that selectively expressed GCaMP6s in excitatory neurons (**Fig. 4c**). In contrast, the frequency of calcium transients and amplitude levels were not different in the prelimbic L5 VGAT-positive inhibitory neurons between control and LSM mice (**Fig. 4d**), indicating that the impact of the microglial N-GAPDH cascade is selectively on the excitatory neurons. These data indicate increased excitation in the prelimbic L5 excitatory neurons of LSM mice, which is normalized by genetic intervention with the N-GAPDH cascade in microglia.

**Fig. 4.**
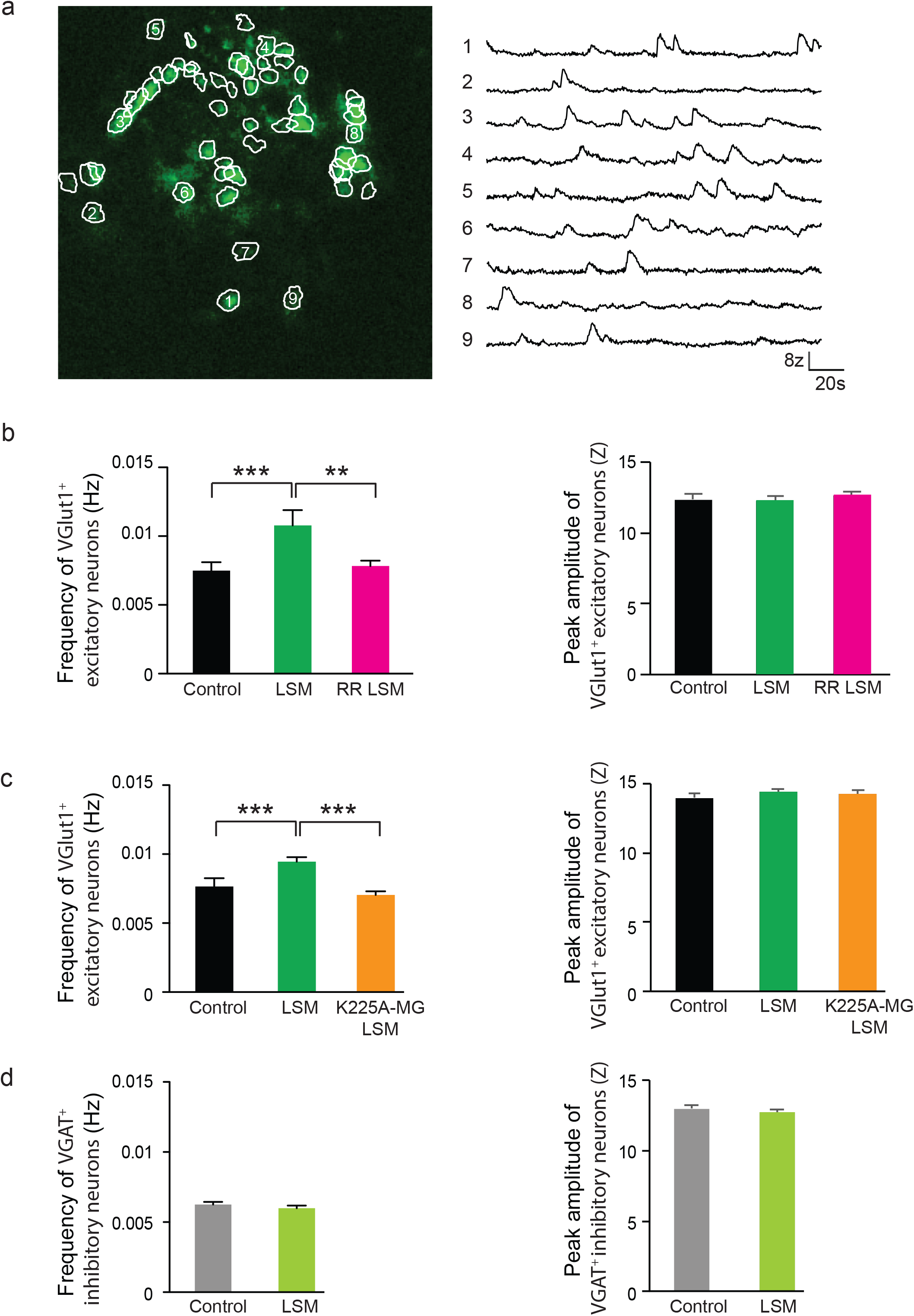
Both pharmacological and microglia-specific genetic intervention to the N-GAPDH cascade normalizes hyperactivation of the prelimbic layer 5 excitatory neurons in LSM mouse model. (**a**) A representative image of a brain slice from a LSM mouse showing expression of VGlut1 neuron specific promoter-driven GCaMP6s in prelimbic cortical neurons (left) and time courses of the fluorescence change (ΔF/F) in the 9 neurons labeled 1-9 in the left panel (right). Brain slices were recorded at basal level and no stimulation was performed. (**b**) Frequency (left) and peak amplitude (right) for 47 VGlut1-positive cells from 4 Control mice, 57 VGlut1-positive cells from 4 LSM mice, and 89 VGlut1-positive cells from 4 RR LSM mice. (**c**) Frequency (left) and peak amplitude (right) for 68 VGlut1-positive cells from 3 Control mice, 238 VGlut1-positive cells from 3 LSM mice, and 171 VGlut1-positive cells from 3 K225A-MG LSM mice. (**d**) Frequency (left) and peak amplitude (right) for 169 VGAT-positive cells from 3 Control mice, and 119 VGAT-positive cells from 5 LSM mice. For each panel, error bars represent the mean ± SEM. **p < 0.01, ***p < 0.001 by one-way ANOVA with Tukey’s multiple comparisons test for panels (**b**) and (**c**). No statistical difference by a two-tailed unpaired Student’s *t*-test for panel (**d**).

## DISCUSSION

The present study provides two major findings of neurobiology and cellular signaling, respectively. First, by using LSM mouse model, we show that stress-induced deficits in rule shifting tasks are mediated by a specific molecular cascade (the N-GAPDH cascade) in microglia, which in turn elicits hyperactivation of the prelimbic excitatory neurons. Previously, the involvement of the neuronal changes in cognitive inflexibility was shown in a genetic model that interferes with neuronal development [3]. In contrast, we now demonstrate a pivotal role of microglia and microglia-neuron crosstalk in stress-induced cognitive inflexibility. Second, we also decipher a mechanism whereby the N-GAPDH cascade mediates stress-induced behavioral alteration, which has been an important question in cellular signaling.

Variety types of stressors elicit cognitive alterations, including deficits of cognitive flexibility such as those in rule shifting tasks. Such stressors may elicit signaling cascades in not only neurons but also non-neuronal cells, such as microglia and astrocytes. The present report offers a prototype of future studies that explore other mechanisms for cognitive alterations elicited by different stressors. In analogy to the microglia-specific genetic intervention selectively disrupting the initial regulatory step for the N-GAPDH cascade that was used in the present study, mouse models that target a specific molecular cascade in a cell type-specific manner will be crucial for future studies.

Here we indicate a causal role of the microglial N-GAPDH cascade in neuronal hyperactivation and behavioral changes. Key questions for the next step, although they are beyond the scope of the present study, will include mechanism(s) that links nuclear translocated GAPDH in microglia to downstream neuronal hyperactivity. Such neuronal changes may be induced by molecules secreted from microglia and/or direct cell-cell interaction between microglia and neurons. Nuclear translocated GAPDH may systematically influence transcriptional landscape in the microglia, given that GAPDH can reportedly interact with DNAs directly or indirectly [40,59]. To address this question experimentally, next generation sequencing at the whole genome level will be useful. Whether and how the N-GAPDH cascade is pathologically manifested in a stress-induced manner in patients with brain disorders may be another outstanding question. To address this question properly, we also need to be sensitive to which cells from patients will be selected for the experiments.

Altogether, we provide a novel conceptual framework that an enzyme, which is traditionally considered for cellular homeostasis through glycolysis, regulates microglial function for adaptive behavior in response to stressors. We speculate that GAPDH is not the only enzyme to have such a unique role. The nuclear localization of other glycolytic enzymes has been historically reported [60,61], however, their role (e.g., transcriptional regulation) remains elusive. We believe that the present study represents the tip of the iceberg for a higher level mechanism that involves the regulation of homeostasis by bioenergetic enzymes, a mechanism that has probably evolved from cellular to complex systems.

## ACKNOWLEDGEMENTS

We would like to thank Hao Zhang and the Flow Cytometry Core at JHSPH for providing sorting and analysis services. We thank Yukiko Lema for figure and manuscript organization. We also appreciate Melissa Landek-Salgado for thoughtful comments and editions of this manuscript. This work was supported by the grants from the National Institute of Health (MH-094268 Silvio O. Conte center, MH-105660, MH-107730), as well as the grants from NARSAD, Stanley, and S-R/RUSK (all to A.S.).

