## Supplementary figures for "Nuclear GAPDH in cortical microglia mediates stress-induced cognitive inflexibility": N-GAPDH bioRxiv Supplementary figures.pdf

### Supplementary Figure 1

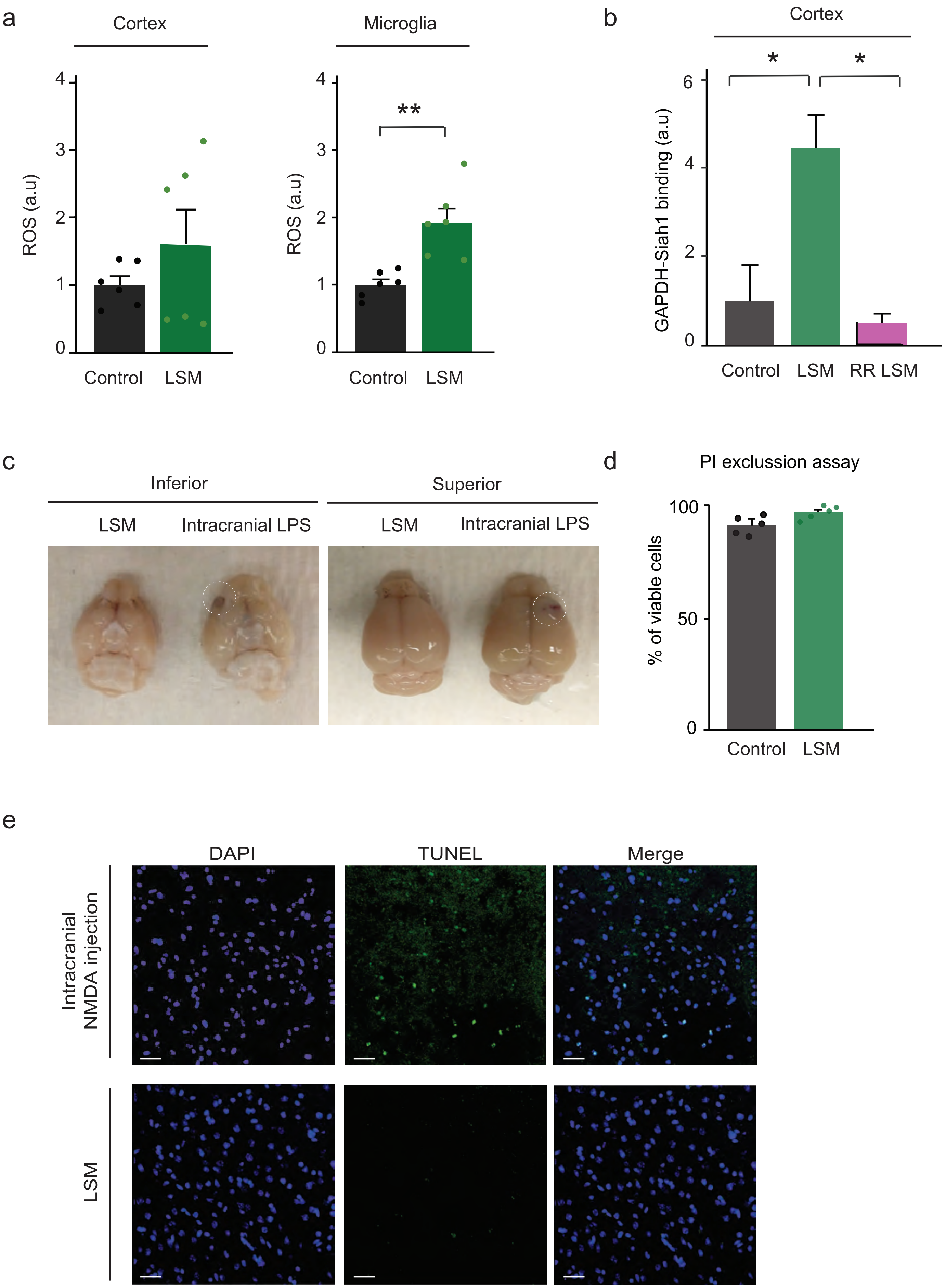

### Supplementary Figure 1

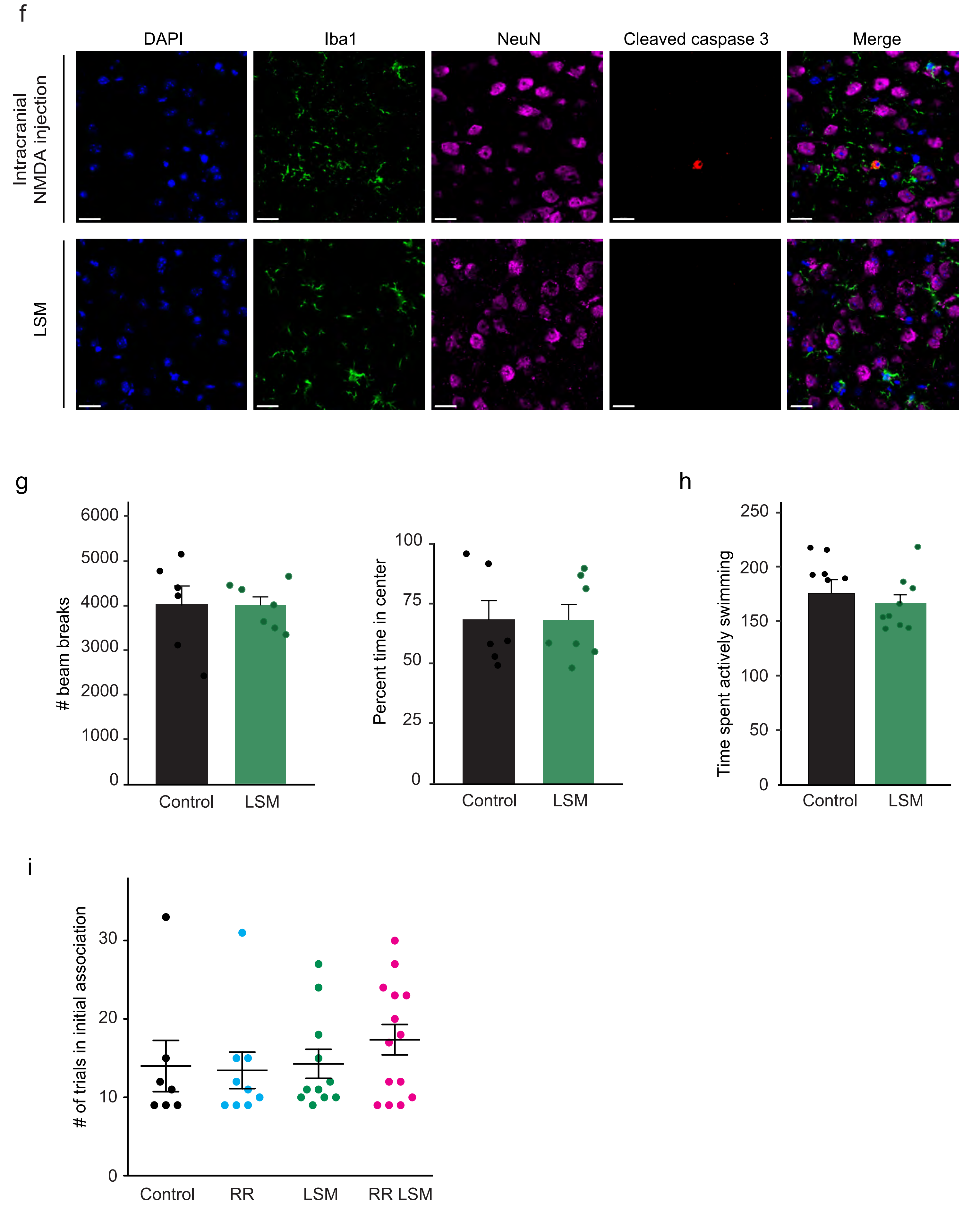

**Supplementary Fig. 1 | Biochemical and Behavioral Characterization of the LSM Model.**

**(a)** The level of reactive oxygen species (ROS) in cortical tissue (left) and cortical microglia (right) from Control and LSM mice measured with the Cell ROX$^{TM}$ reagent for flow cytometry.

**(b)** The level of GAPDH-Siah1 binding, which was normalized by the level of Siah1 in the cortex. Lysates were from cortical brain tissue from Control, LSM, and RR LSM mice.

**(c)** No leakage of Evans Blue in LSM mice. Mice intracranially injected with LPS were used as a positive control. Circles indicate the brain region where Evans Blue leakage was detected in the positive control.

**(d)** Cell death was not detected in cortical tissue from Control and LSM mice. Cell death was assessed by a flow cytometry propidium iodide (PI) exclusion assay.

**(e)** Representative confocal images showing a TUNEL assay for LSM mice. Mice intracranially injected with a 10 μM NMDA solution were used as a positive control. Blue, DAPI (nuclear staining); green, positive signals in the TUNEL assay. Scale bar: 20 μm.

**(f)** Representative confocal images depicting cleaved caspase-3. A cleaved caspase-3 signal was not detected in the brains of LSM mice. Mice intracranially injected with a 10 μM NMDA solution were used as a positive control. Iba1 and NeuN staining were used as cellular markers for microglia and neurons, respectively. Scale bar: 20 μm.

**(g)** Performance of Control and LSM mice during the open field test.

**(h)** Performance of Control and LSM mice during the forced swim test.

**(i)** Performance of Control, RR, LSM, and RR LSM mice in the initial association of the attentional set-shifting task.

Each dot represents data from an individual mouse for panels [**(a)**, **(d)**, **(g)**, **(h)**, and **(i)**]. Bar graph represents data from 3 independent experiments (**b**). Confocal images are representative of 3 mice per group [(**c**), (**e**), and (**f**)]. Error bars represent the geometric mean ± SEM for panels **(a)** and **(d)**, and the mean ± SEM in the other panels. *$p < 0.05$ by one-way ANOVA with Tukey's multiple comparisons test for panels **(b)** and **(d)**; unpaired two-tailed t-test for panels for **(a)**, **(g)**, and **(h)**; and a two-way ANOVA (RR x LPS) test for panel **(i)**.

Supplementary Figure 2

a

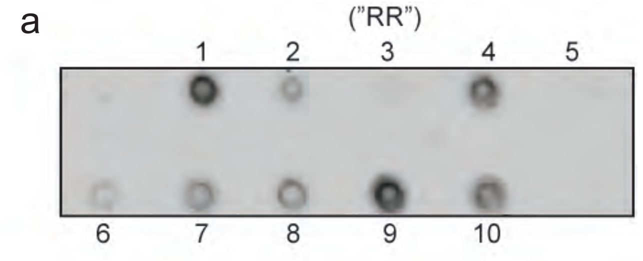


1: No treatment
2: Deprenyl
3: (1R,3R)-1,3-dimethyl-2-propargyl-tetrahydroisoquinoline ("RR")
4: (R,S)-1-methyl-propargyl-tetrahydroisoquinoline
5: (1S,3S)-1,3-dimethyl-2-propargyl-tetrahydroisoquinoline
6: (R,S)-1-methyl-tetrahydroisoquinoline
7: (S)-1-methyl-tetrahydroisoquinoline
8: (R)-1-methyl-propargyl-tetrahydroisoquinoline
9: (1R,3S)-1,3-dimethyl-2-propargyl-tetrahydroisoquinoline
10: (S)-1-methyl-propargyl-tetrahydroisoquinoline

b

| Sample | Serum pmol/mL | Brain pmol/g | Ratio brain/serum |
|---|---|---|---|
| Control | BLQ | BLQ | |
| RR1 | 13.8 | 42.6 | 3.08 |
| RR2 | 10.6 | 28.5 | 2.69 |
| RR3 | 13.2 | 33.3 | 2.52 |
| RR4 | 14.8 | 60.6 | 4.09 |
| RR5 | 16.0 | 42.2 | 2.64 |

**Supplementary Fig. 2 | Characterization of (1R, 3R)-1, 3-dimethyl-2-propargyl-1, 2, 3, 4-tetrahydroisoquinoline "RR."**

**(a)** Dot blot analysis for inhibition of GAPDH-Siah1 binding. (1R, 3R)-1, 3-dimethyl-2-propargyl-1, 2, 3, 4-tetrahydroisoquinoline (the RR compound) blocked the binding between GAPDH and Siah1 the most potently among the listed deprenyl structural analogues.

**(b)** *In vivo* pharmacokinetic analysis of RR demonstrating brain enrichment compared to sera. The control sample was below the limit of quantification (BLQ). RR mice (N=5); Control mouse (N=1).

### Supplementary Figure 3

a

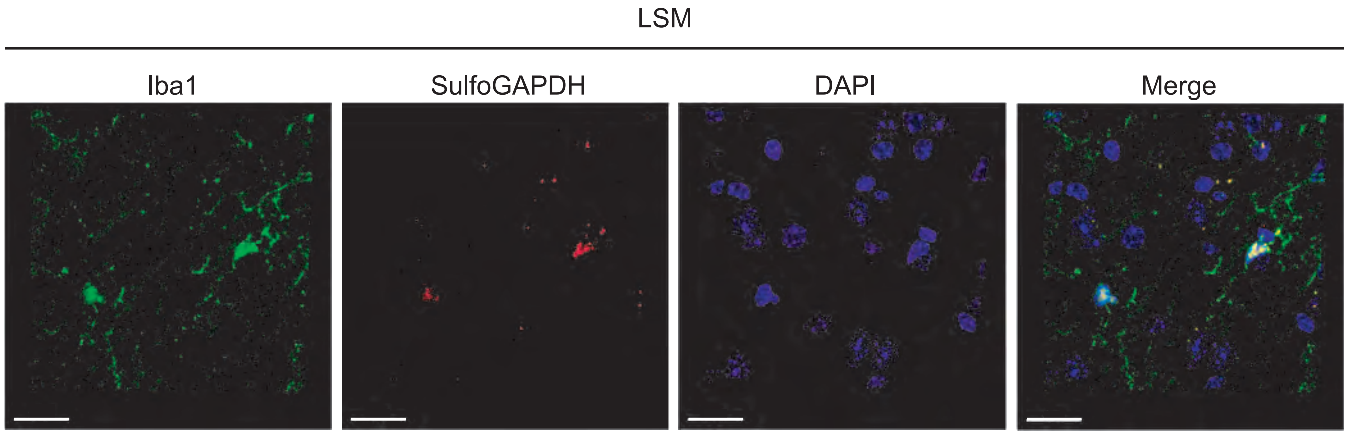


b

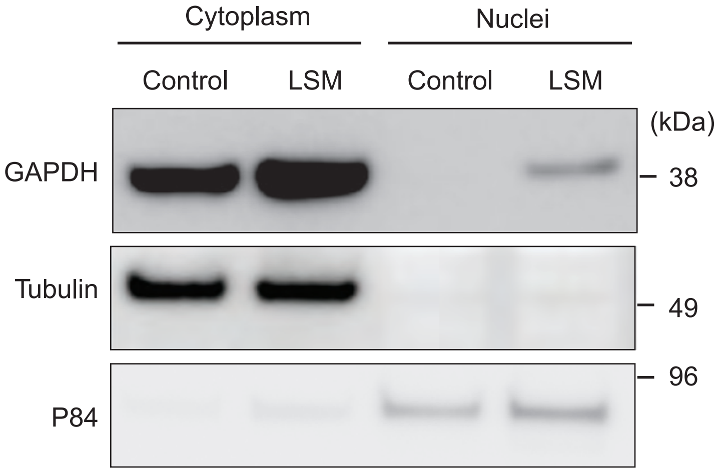


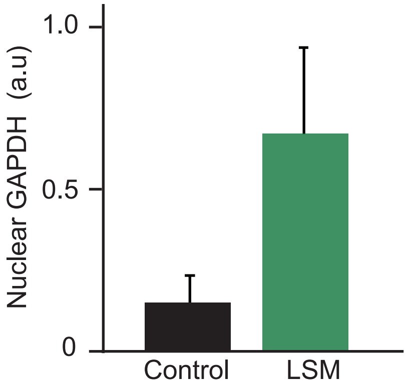

**Supplementary Fig. 3 | Activation of the N-GAPDH Cascade Selectively in Microglia of LSM Mice.**

**(a)** Representative confocal images (63X) for the co-localization of sulfonated GAPDH at Cys-150 (SulfoGAPDH) with the Iba1 microglia-specific marker in LSM mice. Scale bar: 20 μm. Confocal images are representative of 3 mice per group.

**(b)** Immunoblots showing cytoplasmic and nuclear GAPDH in cortical microglia. Immunoblots are representative of 2 independent experiments from nuclear fractionations performed with cortical microglia isolated from 5 mice per group.
